# On Rank Deficiency in Phenotypic Covariance Matrices

**DOI:** 10.1101/2020.07.23.218289

**Authors:** F. Robin O’Keefe, Julie A. Meachen, P. David Polly

## Abstract

This paper is concerned with rank deficiency in phenotypic covariance matrices: first to establish it is a problem by measuring it, and then proposing methods to treat for it. Significant rank deficiency can mislead current measures of whole-shape phenotypic integration, because they rely on eigenvalues of the covariance matrix, and highly rank deficient matrices will have a large percentage of meaningless eigenvalues. This paper has three goals. The first is to examine a typical geometric morphometric data set and establish that its covariance matrix is rank deficient. We employ the concept of information, or Shannon, entropy to demonstrate that a sample of dire wolf jaws is highly rank deficient. The different sources of rank deficiency are identified, and include the Generalized Procrustes analysis itself, use of the correlation matrix, insufficient sample size, and phenotypic covariance. Only the last of these is of biological interest.

Our second goal is to examine a test case where a change in integration is known, allowing us to document how rank deficiency affects two measures of whole shape integration (eigenvalue standard deviation and standardized generalized variance). This test case utilizes the dire wolf data set from Part 1, and introduces another population that is 5000 years older. Modularity models are generated and tested for both populations, showing that one population is more integrated than the other. We demonstrate that eigenvalue variance characterizes the integration change incorrectly, while the standardized generalized variance lacks sensitivity. Both metrics are impacted by the inclusion of many small eigenvalues arising from rank deficiency of the covariance matrix. We propose a modification of the standardized generalized variance, again based on information entropy, that considers only the eigenvalues carrying non-redundant information. We demonstrate that this metric is successful in identifying the integration change in the test case.

The third goal of this paper is to generalize the new metric to the case of arbitrary sample size. This is done by normalizing the new metric to the amount of information present in a permuted covariance matrix. We term the resulting metric the ‘relative dispersion’, and it is sample size corrected. As a proof of concept we us the new metric to compare the dire wolf data set from the first part of this paper to a third data set comprising jaws of *Smilodon fatalis*. We demonstrate that the *Smilodon* jaw is much more integrated than the dire wolf jaw. Finally, this information entropy-based measures of integration allows comparison of whole shape integration in dense semilandmark environments, allowing characterization of the information content of any given shape, a quantity we term ‘latent dispersion’.

## introduction

The study of integration in biological systems has a long history, stretching back to Darwin (1859), given a modern footing by Olson and Miller (1958), quantified by Van Valen (1974) and Cheverud (1982, 1996), and maturing into a broad topic of modern inquiry encompassing genetics, development, and the phenotype (see Klingenberg, 2013, and Goswami and Polly, 2010, for reviews). In a biological context, ‘integration’ means that traits of a whole organism covary to a significant degree. This covariance is critical in evolving populations, because traits are not free to respond to selection without impacting dependent traits, and the directionality of selection response is constrained by these dependencies (Grabowski and Porto, 2017, Figure 1). Yet trait dependency can also remove constraint by giving a population access to novel areas of adaptive space (Goswami et al., 2014, Figure 5). Consequently phenotypic integration is central to the evolvability of biological systems, because it defines the achievable adaptive space.

**Figure 1.**
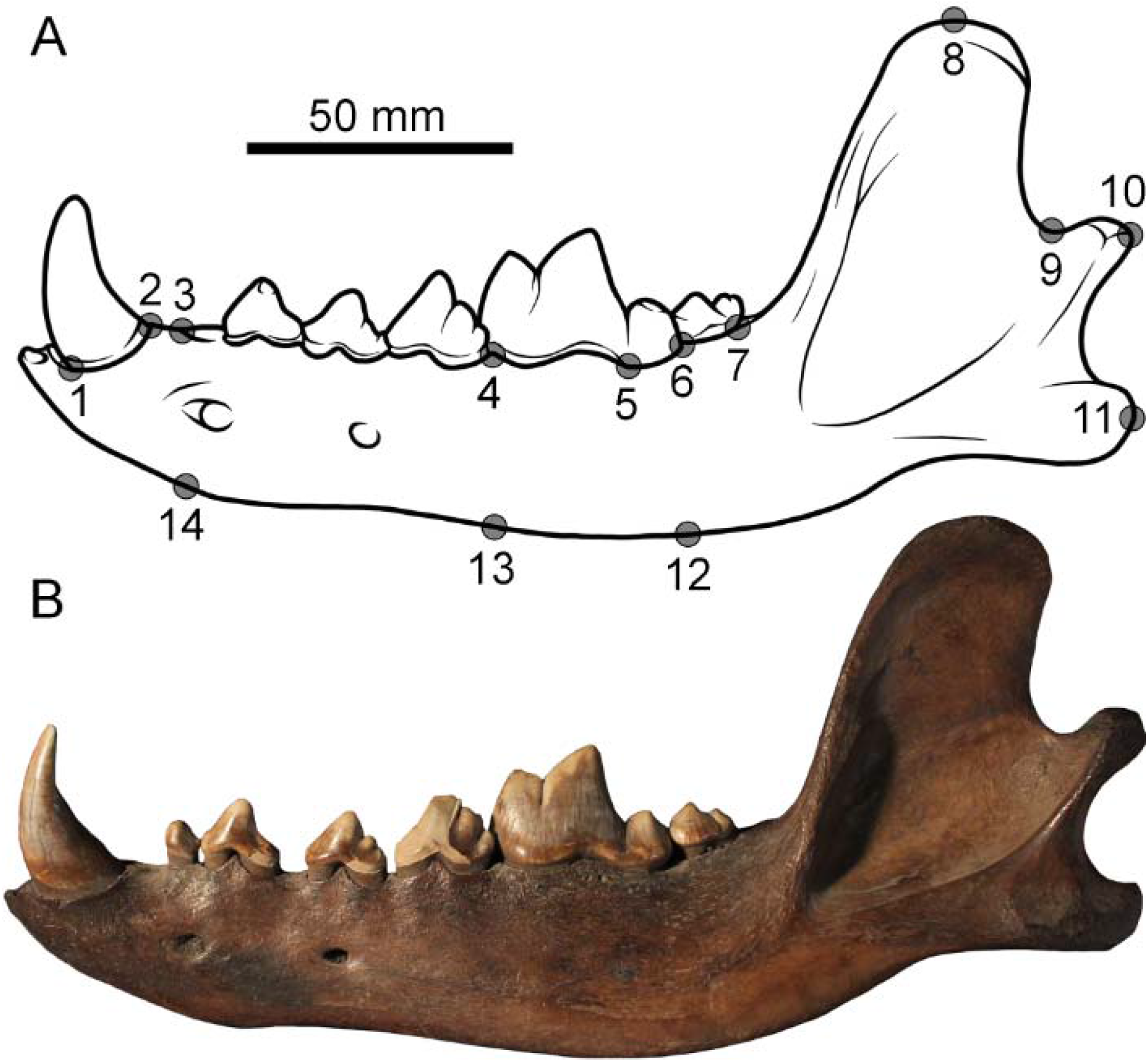
Landmarks used in this study, numbered on a schematic dire wolf jaw (A), and (B) a representative *Canis dirus* dentary from Pit 61/67, Rancho La Brea, California.

### Background

The use of landmark data to characterize biological shape, and to quantify shape change, has become ubiquitous since the introduction of geometric morphometrics by Bookstein (1997; reviewed in Zelditch et al., 2012). Analysis of geometric morphometric landmark data begins with Generalized Procrustes Analysis (GPA), wherein the centroid of each specimen is translated to the origin, and the coordinates are rotated and scaled to a mean shape so that the intra-landmark variance among specimens is minimized (Bookstein 1997). Given a matrix **LM_n,v_** of *v* two- or three-dimensional landmark coordinates and *n* specimens, a matrix resulting from Procrustes superimposition is:

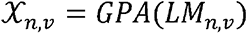

There are myriad examples in the literature of the application of principal components analysis to GPA-transformed landmark data (for example, Segura et al., 2020, for canid dentaries; Brannick et al., 2015, for dire wolf dentaries, and O’Keefe et al. 2014, for dire wolf crania). The usual analysis progression is to calculate the principal component scores for the first few PCs, and plot the specimens into the principal component space. Principal components analysis is therefore a powerful tool for dimensionality reduction, allowing the visualization and biological interpretation of a large proportion of the variance on one or a few axes. Analyses of this type typically utilize the first few eigenvectors, so the magnitude of their associated eigenvalues is not a central concern beyond ensuring their significance. However the eigenvalue vector also carries biological signal, and analysis of its distribution has given rise to the concept of whole-shape phenotypic integration.

### Whole-Shape Integration Measures

> *“The ‘generalized variance’, |Σ*|*, the determinant of the variance-co-variance matrix, is related to the area (or hypervolume) of the equi-probability ellipses (ellipsoids) of the distribution.”*—VanValen, 1974, p. 235.

The distribution of the eigenvalue vector has formed the basis of several metrics intended to quantify the strength of covariation in a whole shape. As stated by VanValen in the quote above, the shape space containing the objects of interest can be conceptualized as a hyperellipse in the original variable space. Principal components analysis moves this hyperellipse to the origin, rotates it, and defines its principal axes. VanValen realized that this hyperellipse had two salient properties: 1) its dimensionality, and 2) its variance, or dispersion, on each axis, which he called ‘tightness’. In subsequent work the dispersion of the eigenvalues of a phenotypic covariance matrix—this tightness— has received a great deal of attention, while its dimensionality has not (reviewed in Najarzadeh, 2019). Cheverud (1983) introduced the first modern measure of trait integration, defining it as

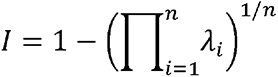

or the geometric mean of the eigenvalues of the correlation matrix subtracted from unity. This quantity is identical to one minus the *n*th root of the generalized variance (Wilks, 1932) proposed by VanValen in 1974. In a wider statistical context this quantity is termed the ‘standardized generalized variance’, or SGV (SenGupta, 1987). The SGV is the geometric mean of the eigenvalue variance, and so may be thought of as the mean diameter of the axes in the hyperellipse. Both Cheverud (1983) and SenGupta (1987) state that the SGV is comparable among spaces of different dimensionality, and this assumption is widely accepted, although it has never been tested (e.g. Najarzadeh, 2019). While both the Cheverud integration and the equivalent SVG have been used as measures of dispersion, a consensus has emerged that the standard deviation of the eigenvalues is a better measure statistically (Pavlicev et al., 2009). Several other metrics have been proposed for quantifying the dispersion of phenotypic matrices, summarized by Pavlicev et al., 2009, while the sample size requirements of various metrics are evaluated by Grabowski and Porto, 2017.

This paper explores the performance of the eigenvalue dispersion and the SGV in rank-deficient geometric morphometric matrices, and we define these metrics here. Eigenvalue dispersion is measured as the standard deviation of the eigenvalue vector of the correlation matrix of GPA landmarks, standardized to the mean eigenvector (Goswami and Polly, 2010, modified from Equations 7 and 8):

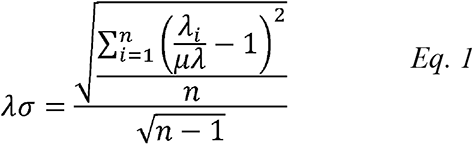

In geometric morphometric data sets, either four (2D) or five(3D) degrees of freedom are lost to the Procrustes analysis, so the number of non-zero eigenvalues will be reduced by this amount (Zelditch et al., 2012). This will also be true of the SGV, calculated as the n^th^ root of the first n-4 eigenvectors of the correlation matrix. Because we use two-dimensional data sets with 14 landmarks in this paper, the correct formulation of the SVG is:

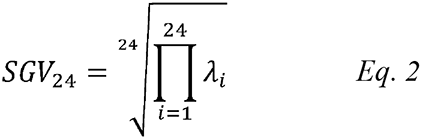

### Integration and Modularity

> *“However, if the structure of a dataset is strongly modular, with several different groups of strongly co-varying traits, then variance will be distributed more evenly across a number of principal components, and eigenvalue variance will be relatively low. Thus, the dispersion of eigenvalues provides a simple measure for comparison of the relative integration or modularity of the structures described in a matrix.”*—Goswami and Polly, 2010, p. 226.

In the 21^st^ century the study of GPA covariance matrices has grown to include the field of modularity and integration (Goswami et al. 2014; Klingenberg, 2013). This conception of integration is similar to Van Valen’s and Cheverud’s in that it studies covariance structure, but differs in attempting to identify variable subsets that covary relative to others. Rather than relying on a single overall measure of dispersion, modularity studies test specific models of landmark covariation against a null model in an attempt to characterize subsets with high intraset connectivity and low interset connectivity (Goswami and Polly, 2010). Because these sets are primary covariance structures in the data, they will exist on the first several principal components, and the entire eigenvalue distribution is not directly relevant. This approach to modularity has yielded powerful hypotheses of evolutionary plasticity and constraint linking development to phenotype, and to evolutionary changes in modularity over deep time (Goswami et al., 2015). Modularity models were first assessed using the RV coefficient introduced by Klingenberg (2008). This coefficient has been superseded by the similar covariance ratio statistic (CR, Adams, 2016), which is more robust to differences in sample size and data dimensionality. Another common approach to assessing modularity model significance is Partial Least Squares (PLS; Goswami and Polly, 2010); this approach is more involved mathematically but has the same goal as the CR statistic (for a recent example on dog and wolf crania see Curth et al., 2017). As stated in the quote above, the modularity concept can also be applied to a whole shape. In whole-shape modularity, a shape would be more integrated if it had greater eigenvalue dispersion, and would be more modular if it possessed lesser dispersion. This is a specific prediction: in a population where modularity is evolving, an increase in modularity should lead to a decrease in integration and hence dispersion, and thus a decrease in dispersion metrics like SVG and eigenvalue standard deviation.

### Rank Deficiency in Phenotypic Covariance Matrices

> *It [PCA] is used to obtain a more economic description of the N-dimensional dispersion of the original data by a smaller number of “principal components”, which are formally the eigenvectors of the dispersion matrix… Hence the number of principal components that contribute significantly to the variation of the sample is the actual “dimensionality” of the dispersion.* – G. P. Wagner, 1984, pp. 92-93.

The rank of a matrix is the dimensionality of the vector space spanned by its variables. If all variables are completely independent then the matrix is of “full rank,” equal to the number of variables, but when one variable depends completely on another, the matrix has “deficient rank”. The full rank value of a matrix is the number of columns, while the “real rank” is the maximum number of linearly independent columns. The derivation of the covariance or correlation matrix **K** directly from **X**,

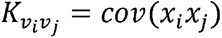

often results in a covariance matrix that is overdetermined; the high degree of correlation in biological systems often results in matrices whose full rank is much higher than the real rank (Van Valen, 1974; Adams, 2016). Significant rank deficiency in **K** will impact the eigenvalue distribution, because eigenvalue distributions of rank deficient matrices will have tails of very small non-zero eigenvalues. This tail of small eigenvalues may impact integration metrics that rely on the eigenvalue distribution, but this subject has not been treated explicitly. Characterizing this problem of rank deficiency, and developing treatments for it, are the goals of this paper.

In geometric morphometric covariance matrices, rank deficiency can arise from at least three sources. These are the Generalized Procrustes Analysis itself, lack of matrix information, and phenotypic covariance. Of these, only phenotypic covariance is of direct biological interest, while the first two may confound attempts to measure it. The magnitude of rank deficiency due to Procrustes analysis is known; the translation, scaling, and rotation performed during the analysis remove four degrees of freedom from two-dimensional data (Zelditch et al., 2012). A covariance matrix derived from **X** should therefore possess *v* – 4 non-zero eigenvalues, or in other words be deficient by four ranks from the full rank.

The rank deficiency resulting from the information intrinsic to **X** is harder to quantify. This lack of information can have two sources: insufficient sample size and landmark oversampling. Lack of information due to sample size arises from too few sampled specimens. Grabowski and Porto (2017) studied the sample size requirements for a range of integration metrics and concluded that an *n* of greater than 100 was required to achieve values stable to changes in *n*. Integration studies with sample sizes of less than 100 are therefore information poor, and this should increase rank deficiency. What we term landmark oversampling refers to the dense landmarking of a relatively simple shape. A simple shape with few modes of variation requires few landmarks to characterize fully (a circle requires one landmark). Use of more than the necessary number of landmarks will result in covariance among the variables, and increased rank deficiency. This source of rank deficiency has also not been quantified. However it is tractable, as we outline in the Discussion.

Another factor impacting matrix rank is use of the correlation matrix rather than the covariance matrix. Use of the correlation matrix artificially inflates matrix rank, because each coordinate of each variable is awarded a full rank. This tacitly assumes that the coordinates within a landmark are uncorrelated, and that variance magnitudes in each coordinate direction are equivalent. However landmark displacements that are dominated by a vector parallel to one axis will have trivial variation on the other, and in a correlation matrix this trivial variation will be awarded a full rank and expanded to a variance of one. A correlation matrix of **X** will therefore possess a degree of noise from inflation of trivial variance (for discussion of the desirability of the covariance matrix in morphometric applications see Goswami and Polly, 2010, p. 217). Therefore use of the correlation matrix is unwise with morphometric data unless some procedure is used to concatenate the landmark coordinates into a single measure before analysis (Goswami and Polly, 2010, and previous authors have used the congruence coefficient, although this has been criticized by Klingenberg 2008).

### Information Entropy and Effective Rank

The first to point out the problem of redundancy in the full and real ranks of **K** in a morphometric context was Van Valen (1974), who tailored an information metric to correct for it. The problem of rank deficiency can be framed in terms of the eigenvalues of a matrix, with the true rank represented by the subset of eigenvalues carrying significant variance. This differs from the set of eigenvalues that carry non-zero variance, which is how matrix rank is usually defined. The question of the number of ‘significant’ eigenvalues is a general question in data analysis and there are many techniques for determining this number, summarized by Cangelosi and Goriely (2007) in the context of cDNA microarray data. They include the broken stick model, Cattell’s SCREE test, and Bartlett’s sphericity test among many others (see also Hine and Blows, 2006; Bunea et al., 2011). All these techniques share fundamental shortcomings, such as an integer value and recourse to an *ad hoc* criterion to assess the cutoff for significance. Yet the problem of redundancy in eigenvalue distributions is general, and has been solved in signal processing contexts by the use of “information entropy” (Shannon, 1948). Information entropy is a metric used to represent the information content of a system by determining how much a signal can be compressed without loss of information. At least two metrics based on Shannon’s information entropy have been proposed to characterize the dimensionality of covariance matrices in a biological context (for a thorough mathematical development see Cangelosi and Goriely 2007). Those authors develop a dimensionality metric for cDNA microarray data they termed the ‘information dimension’. A very similar metric was developed by Roy and Vetterli (2007); those authors term their metric the ‘effective rank’, and use it synonymously with ‘effective dimensionality’. The following application of the Shannon entropy to geometric morphometrics follows the Roy and Vetterli (2007) development of effective rank.

Given a GPA covariance matrix **K** with eigenvalues Λ**_V_**:

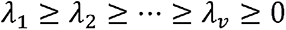

The Shannon entropy *E_S_* of **K** is defined as

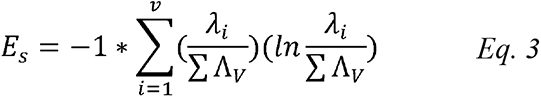

Note that the eigenvalues are standardized to the trace of Λ; this is necessary because the information entropy was originally defined in a probability context, and the terms being evaluated must therefore sum to unity (Shannon, 1948). Roy and Vetterli define their effective rank as *e* raised to the power *E_s_*; the ‘effective rank’ of **K** is defined in the same way:

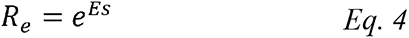

This effective rank, *R_e_*, of **K** is a continuous metric, is based on solid theoretical grounds from information theory, and may be thought of as the signficant number of dimensions of the shape space represented by **K**, or equivalently as the numner of non-redundant eigenvalues in **K**.

### Study Design

This paper has three main components, with the later dependent on the results of the earlier. Therefore we divide the Materials and Methods section into an independent Materials section, followed by three sections of Methods and Results. This organization allows sequential presentation of results as they become necessary for further methodological development. The following sections are:

#### Materials

This section introduces the three data sets used in this paper. These comprise two samples of dire wolf jaws from Rancho La Brea (RLB), originating from tar pits deposited about 5000 years apart. The third data set consists of jaws from the sabertooth cat, *Smilodon fatalis*, assembled from multiple RLB pits (*n* = 81). All three data sets comprise 14 two-dimensional landmarks taken from photographs of jaws in lateral view.

#### Methods and Results, Part 1

In this section, we use the concept of effective rank to investigate the rank deficiency of the Pit 61/67 dire wolf population. We demonstrate that the covariance matrix based on these data is highly rank deficient. Through use of a permutation test, we characterize the magnitude of the rank deficiency arising from lack of matrix information. When combined with the rank deficiency expected from the GPA, this allows us to quantify the rank deficiency due to phenotypic covariance. Lastly, we quantify the rank inflation arising from use of the correlation matrix.

#### Methods and Results, Part 2

Here we evaluate the impact of the observed rank deficiency on whole shape integration metrics. We do this by introducing the Pit 13 dire wolf sample, and demonstrating that it is more integrated than the Pit 61/67 sample. This requires testing each population for candidate modularity models, and demonstrating that Pit 61/67 wolves are significantly modular (best described by a two-parameter model), while the Pit 13 wolves are more integrated (best described by a one-parameter model). Current whole shape integration metrics fail to capture this change in integration; therefore we develop a modified version of the SVG based on the effective rank. Because the Pit 61/67 sample size is larger than Pit 13, we can jackknife the Pit 61/67 sample size down to that of Pit 13. This jackknife yields both a confidence interval for hypothesis testing, and normalizes the rank degeneracy due to matrix information content, allowing us to demonstrate that the new metric is successful in capturing the increase in integration in Pit 13 wolves. We term this metric the ‘effective dispersion’.

#### Methods and Results, Part 3

In this section, we generalize the concept of effective dispersion to the case of arbitrary matrix information content. The general metric developed here, termed the ‘relative dispersion’, allows comparison of whole shape integration among data sets of different sample sizes. It also allows for comparison among data sets of arbitrary landmark number, because it is normalized to the information content of the input variables. This normalization relies on the computation of the effective rank of a permuted data matrix. We demonstrate the utility of the relative dispersion metric by analyzing the *Smilodon* data set, finding that the *Smilodon* jaw is much more tightly integrated than that of *Canis dirus*.

The last analytical section is followed by a standard Discussion. One topic considered is the extension of the relative dispersion into a dense semilandmark context, allowing quantification of the information content of a comnplete shape. We propose the term ‘latent dispersion’ for this quantity.

## MATERIALS

### Background

The primary data in this paper come from two populations of dire wolves (*Canis dirus*, Figure 1) from the Rancho La Brea tar pit lagerstätten, deposited in what is now Los Angeles during the terminal Pleistocene (Stock and Harris, 1992). The populations come from two pit deposits, Pits 13 and 61/67. The Pit 13 wolves show elevated tooth breakage and wear, while those from 61/67 do not (Binder et al., 2002). Elevated tooth breakage and wear are thought to indicate nutrient stress, as animals both process carcasses more completely, and compete more intensely, when prey resources are scarce (Van Valkenburgh, 1988, 2009; Meloro, 2012). Subsequent work established this pattern beyond doubt and was able to assign approximate ages to the relevant pit deposits (Binder and Van Valkenburgh, 2010; O’Keefe et al., 2014; Brannick et al., 2015). Phenotypic differences between pits have also been documented; morphometric analyses of dire wolf jaws (Brannick et al., 2015) and crania (O’Keefe et al., 2014) detected clear phenotypic differences in Pit 13 wolves relative to those in 61/67. Pit 13 wolves are significantly smaller in overall body size, and also show significant shape differences, including diminution of the viscerocranium relative to the neurocranium. The observed size and shape changes were attributed to a chronic lack of nutrition during development, producing a neotenic adult phenotype in Pit 13. The biological drivers behind this evolutionary change are not relevant here; what is relevant is that there is a significant modularity shift between the two pits. This creates a useful model system comprising two populations of the same species deposited in the same place, separated only in time, and possessing a significant change in modularity.

### Data Collection

Fossils utilized in this paper are left dentaries of *Canis dirus*, from Pits 13 and 61/67, housed in the Tar Pit Museum at Hancock Park, Los Angeles. The input data for all analyses were 14 landmarks taken from 119 dire wolf jaws, a subset of the 16 landmark data set from Brannick et al. (2015). One hundred and nineteen adult *Canis dirus* dentaries were included in this study. The labial side of specimens from Pits 13 (n =36), and 61/67 (n = 83) were photographed using a tripod-mounted Cannon EOS 30D 8.20-megapixel camera. All specimens were anatomical lefts and were laid flat and photographed with a 5 cm scale-bar. While camera angle and distance were held constant when photographing, scale-bars were used to properly size specimen images in Adobe Photoshop CS2 v.9.0 before landmark digitization; scaling was mostly unnecessary, and always minor.

Fourteen homologous landmarks were digitized on each specimen using the program tpsDig2 (Rohlf, 2013). Positions of landmarks were chosen to give a general outline of the mandible and capture information of functional relevance (Brannick et al., 2015; Figure 1). Landmarks on the tooth row were placed on the alveolus so specimens with missing teeth could be included. However, presence of the lower carnassial tooth was required in order to obtain landmark 5. Two landmarks from Brannick et al. were not used; one of these, at the angle of the mandible, was difficult to identify and showed unacceptably large variance relative to the other landmarks. The second was an interior landmark in the masseteric fossa; it was deleted to make the dataset amenable to outline-based iterative semilandmarking.

Lastly, as a demonstration of the utility of the relative dispersion metric in Part 3, we analyze a data set of *Smilodon fatalis* jaws (Meachen et al., 2014), again from Rancho La Brea. The data set comprised 14 landmarks digitized from photographs of 81 *Smilodon* jaws originating from Pits 61/67 and 13 at the Tar Pit Museum, and the UCMP collection. Pooling of the *Smilodon* sample was necessary to increase sample size.

## MATERIALS AND RESULTS

### Part 1: Rank Deficiency in Pit 61/67

In this section, we examine the amount of rank deficiency of covariance matrices derived from the Pit 61/67 *Canis dirus* data set, and quantify where it is coming from. Standard Procrustes superimposition of the landmark data was first performed using the **geomorph** package (Adams et al., 2020) in R (R Core Team, 2014). Procrustes superimposition yielded a matrix of transformed (GPA) landmarks **X** and a vector of centroid size. Principal components analysis of **X** was performed on four covariance matrices. These were the correlation and covariance matrices for the original data, and correlation and covariance matrices for permuted data. The scree plots for the eigenvalue vectors of the four matrices are shown in Figure 2. Also shown in Figure 2 are the effective ranks of the four matrices, calculated as defined in Equations 3 and 4. Code for computing the effective rank of covariance and correlation matrices, including the permutation code, was written in R (R Core Team, 2014; see Appendix 1). The effective ranks calculated from the correlation and covariance matrices derived from the real data are unique values. Effective rank was also calculated for 10000 matrices in which each column was permuted; this permuted mean is plotted and the standard deviations are reported in the caption to Figure 2.

**Figure 2.**
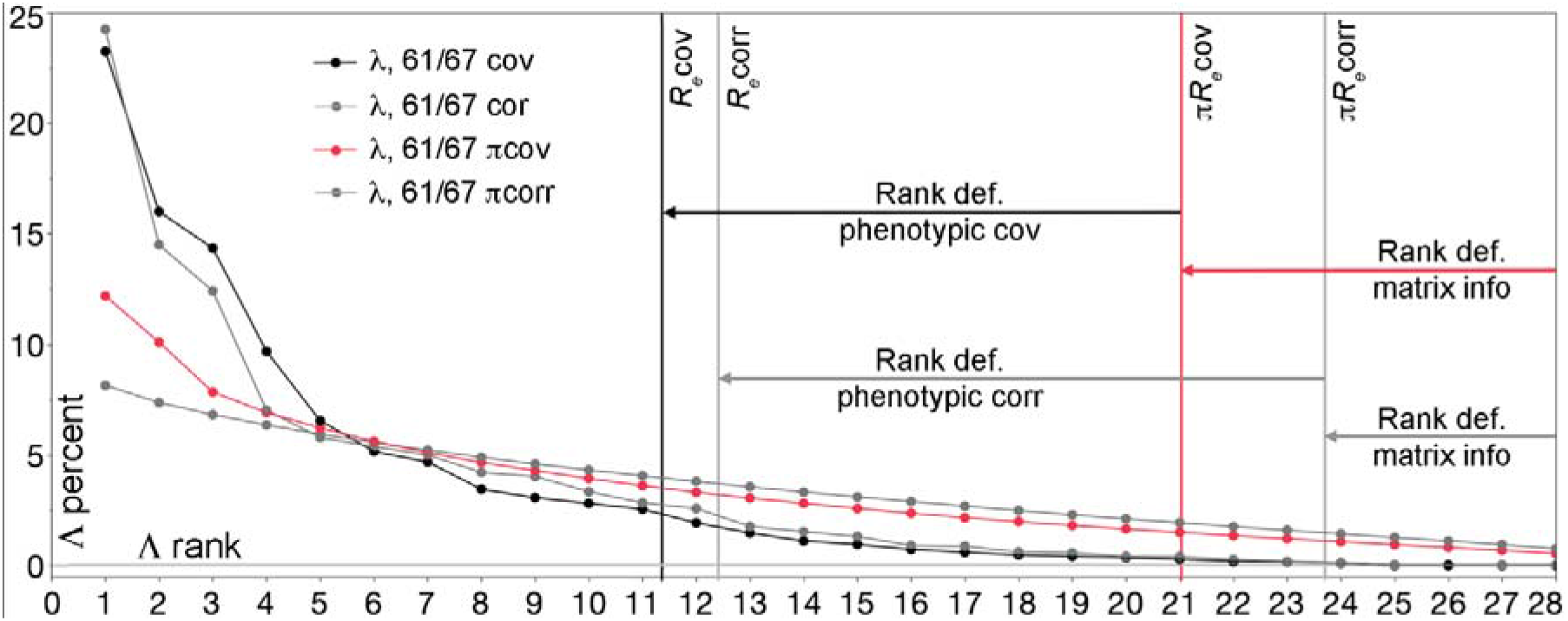
Rank deficiency and its sources in Pit 61/67 dire wolves. Scree plots for four data treatments are plotted. Note that the last four eigenvalues are exactly zero for both the correlation and covariance matrices computed from the real data, reflecting the loss of four degrees of freedom to the GPA. The effective rank of permuted data shows rank deficiency due to lack of matrix information; the true effective ranks are much smaller than this, and this additional deficiency is due to phenotypic covariance. Effective ranks for each matrix are: Permuted covariance = 21.028 +/− 0.226; permuted correlation = 23.728 +/− 0.258; covariance = 11.362; correlation = 12.424.

The scree plots of both the correlation and covariance matrices of X demonstrate that the vector of eigenvalues for each matrix contains 24 non-zero eigenvalues, the expected number given the loss of four ranks to the GPA. Eigenvalues 25-28 are non-zero in the permuted matrices, showing that this rank deficiency is recovered when the matrix is randomized. Effective rank calculations show that both matrices are also highly rank deficient. The rank of the covariance matrix is 11.362, meaning that less than half of the 24 non-zero eigenvalues carry meaningful information. The number of ranks in the permuted covariance matrix is about 21, indicating that the covariance matrix is information poor by about seven ranks. Four of these are from the GPA; the rest are probably due to insufficient sample size (Grabowski and Porto, 2017). The rank deficiency due to phenotypic covariance is therefore 21 minus the effective rank of the real matrix (11.36), or about 9.64 ranks. Therefore 12.64 ranks, or over half, of the eigenvalue distribution is redundant. The small values of the higher eigenvalues are caused mainly by phenotypic covariance, but they do not carry information about it, and lack of matrix information acerbates this deficiency. The eigenvalue vector is overdetermined with respect to its information, and its distribution will be affected as a result.

Figure 2 also shows that the rank of the correlation matrix is 12.424. Use of the correlation matrix therefore adds over one full rank of information relative to the covariance matrix. We believe this rank increase is noise, as described in the Introduction. The pattern for the permuted matrices is similar; the correlation matrix adds almost three ranks of noise over the value for the covariance matrix.

### Part 2: Modularity Models and Whole Shape Integration Metrics

In this section, we introduce the Pit 13 dire wolves and show they are more integrated than those from Pit 61/67. We do this by testing both populations for modularity against a range of candidate two- and three parameter models. We then calculate the eigenvalue distribution and SVG for the two covariance matrices, and show that both fail to capture the modularity change. We then modify the SVG to use only the non-redundant eigenvalues, and show that this measure does capture the modularity change.

#### Modularity Models

A challenge in all modularity studies is initial specification of the models to be tested. In this paper we use a heuristic approach: examining the multivariate allometry vectors of interlandmark distances to identify candidate models. While important, the specification of models is tangential to our discussion of rank deficiency, and we include it below as Appendix 2. This allows us to move directly to testing the candidate models for the two data sets. A total of 14 models of modularity were suggested by the multivariate allometry analyses of the interlandmark distances, and these models were tested individually using the CR statistic (Goswami and Polly, 2010; Figure 3; Table 1), performed in the **geomorph** package in R (Adams et al., 2020). The models were tested both on the pooled data, and on data divided by pit. Effect-size tests and tests against a model of zero modules were performed on 61/67 wolves only, as only these wolves displayed significant modularity; representative tests and results are listed in Table 1.

**Figure 3.**
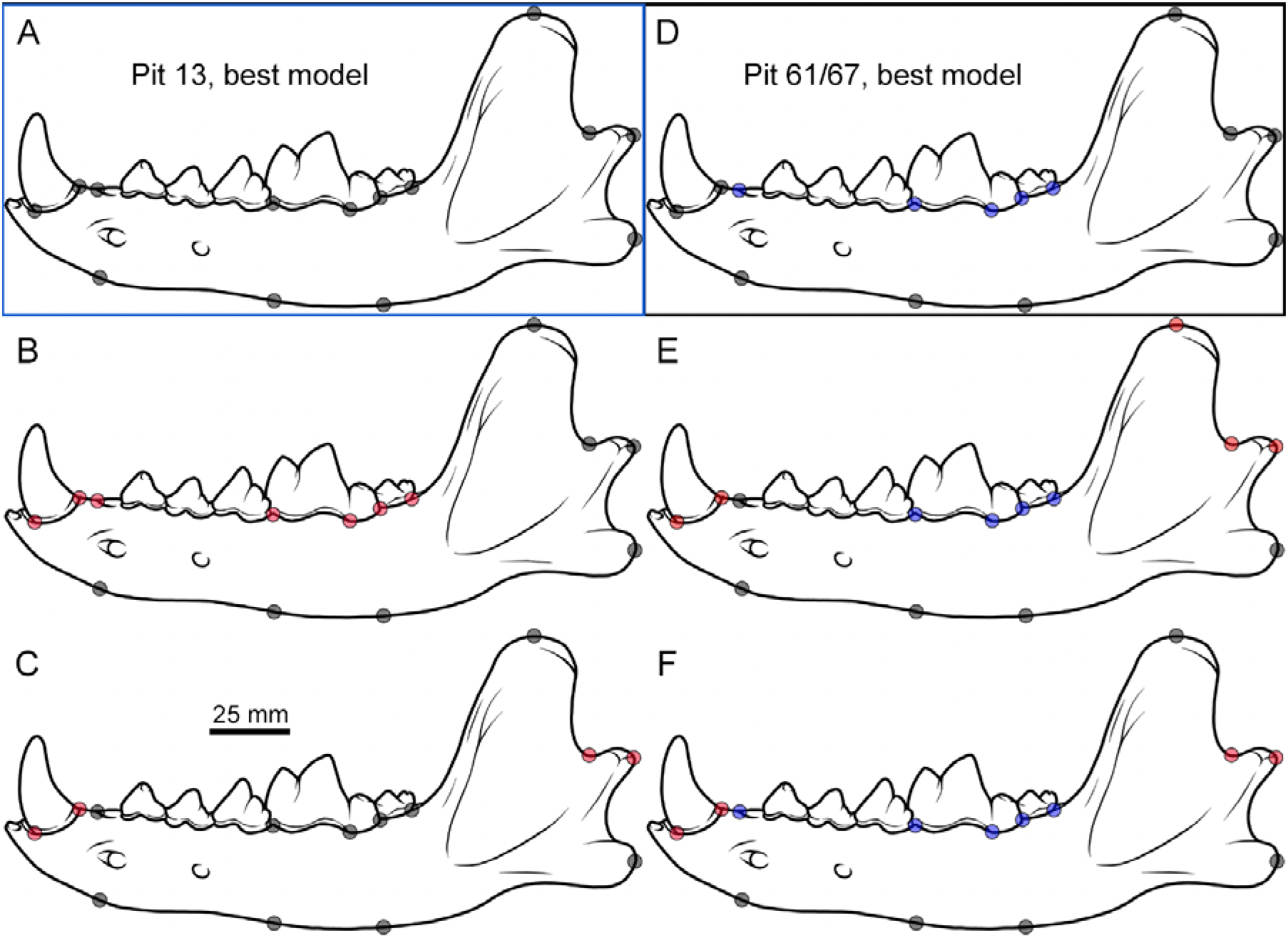
Candidate modularity models for the *Canis dirus* jaw tested in this paper. The preferred modularity model for Pit 61/67 wolves is D, a two-parameter model of (3,4,5,6,7) vs. all other landmarks. This is the model with the best statistical support, but only in Pit 61/67 (Table 1). The preferred model for Pit 13 wolves is A, the one-parameter model. The other models shown did not have significant modularity according to the CR statistic; all models tested are listed in the caption to Table 1.

**Table 1.** Results of representative modularity model tests. Representative module models are listed on the left column, with CR statistics for three data partitions in subsequent columns. Pit 61/67 wolves have two significant modules comprising the check teeth vs. the rest of the jaw, while Pit 13 wolves do not show significant modularity between these or any other modules. As shown in Figure 3, Pit 13 wolves are best fit with a one-parameter model, while Pit 61/67 wolves are best fit with a two-parameter model. Wolves from 61/67 are therefore significantly more modular than those from Pit 13. Other modularity models tested that were not significant include the two-parameter models (4,5,6,7,13,14), (4,5,6,7,8), (4,5,6,7,8,13,14), (1,2,8,9,10), and (1,2,14,9,10,11), and the three-parameter models ((1,2,8,9,10)(4567)), ((1,2,9,10)(4,5,6,7,13,14)), and ((129,10)(4,5,6,7,8)).

| Model, vs. all<br>other LMs | CR<sub>pooled</sub> | P value | CR<sub>13</sub> | P value | CR<sub>61/67</sub> | P value | P<sub>null</sub> |
| --- | --- | --- | --- | --- | --- | --- | --- |
| 1,2 | 1.17 | 0.427 | 1.234 | 0.581 | 1.063 | 0.219 | - |
| 1,2,9,10 | 1.10 | 0.573 | 1.201 | 0.838 | 1.090 | 0.491 | - |
| 1-7 | 1.15 | 0.837 | 1.147 | 0.786 | 1.171 | 0.921 | - |
| 3-7 | 0.96 | 0.052~ | 1.067 | 0.387 | 0.898 | 0.022* | 0.031* |
| 4-7 | 0.92 | 0.046* | 0.994 | 0.108 | 0.894 | 0.028* | 0.046* |
| 1,2,9,10;<br>3,4,5,6,7 | 1.06 | 0.244 | 1.192 | 0.719 | 1.029 | 0.226 | - |

There are two models with statistically significant CR values (Table 1). Both are two parameter models that contrast the length of the post-canine tooth row with the rest of the jaw. The effect sizes of the modularity models with significant CRs were compared using the **compare.CR** function in **geomorph**; the effect sizes were not significantly different. Comparison with a null model of zero modules was performed using the same function. For Pit 61/67 wolves only, the cheek tooth model (3-7) was significantly better than the null model, while the model containing only the molars (4-7) was marginally so. The anterior-posterior width of the canine and of the condyloid process (1,2,9,10) do not form a distinct module on their own; the significant CR of the three-module model is driven by the presence of the cheek tooth module. The cheek teeth are clearly a module. The canine does not take part in this module, and therefore has significant variance attributable to another factor. The inability of the modularity tests to identify the canine and condyloid process as a module was surprising. We believe it was not identified because most of its variation covaries with size. The cheek teeth vary against size, and this allows identification of that module. We note that Pit 13 wolves had no significant modules, and must therefore be more integrated than Pit 61/67 wolves.

The magnitude of the CR differences between pits was surprising, given they are taken from populations of the same species at the same location, sampled 5000 years apart. Given that the sample size of Pit 13 was 36 while that of 61/67 was 83, we were concerned that a lack of statistical power was obscuring the modularity signal in Pit 13. To test for this, we jackknifed the 61/67 sample to 36 members to mirror the sample size of Pit 13 before re-running the CR analysis. Although confidence intervals widened in these subsets, the covariance structure of Pit 61/67 was still clearly evident and significantly modular, indicating that the modularity difference between pits is not attributable to difference in sample size. The pooled CR statistics, run on all 119 wolves, indicate that the inclusion of the Pit 13 wolves actually degrades the global modularity signal (Table 1). This indicates that the relative lack of modularity in Pit 13 is real.

#### Measures of Whole-Jaw Integration

the eigenvalue dispersion and SVG metrics were calculated using eigenanalysis, bootstrapping, and permutation codes written in R (R Core Team, 2014; Appendix 1). The eigenvalue standard deviation metric is significantly less in Pit 13 wolves (Table 2: *λσ*, 13 < 61/67, *p* = 0.00008), while the SGV_24_ metric is equivalent between the two groups (13 = 61/67, *p* = 0.837, Student’s *T* in this and subsequent comparisons). This implies that eigenvalue dispersion is significantly less in Pit 13, and those wolves should be more modular than Pit 61/67. Yet this is not the case; the modularity tests reported above clearly show that Pit 61/67 wolves are significantly modular, while Pit 13 wolves are not. The eigenvalue dispersion is less in Pit 13 even though the jaws are more integrated. The SGV_24_ metric fails to detect the increase in modularity. As demonstrated above, over half of the ranks used to calculate both metrics are redundant, and we believe the SGV_24_ is essentially measuring noise. We experimented with the use of the covariance matrix in the SGV_24_ to see if the failure of that metric was due to use of the correlation matrix. Use of the covariance matrix required standardization of the eigenvalues before calculation to control for different matrix variances. This modified metric is also reported in Table 2, and like the SGV_24_ it is equivalent between the two matrices. Rank inflation due to use of the correlation matrix is therefore not the reason for failure of the SGV_24_.

**Table 2.**
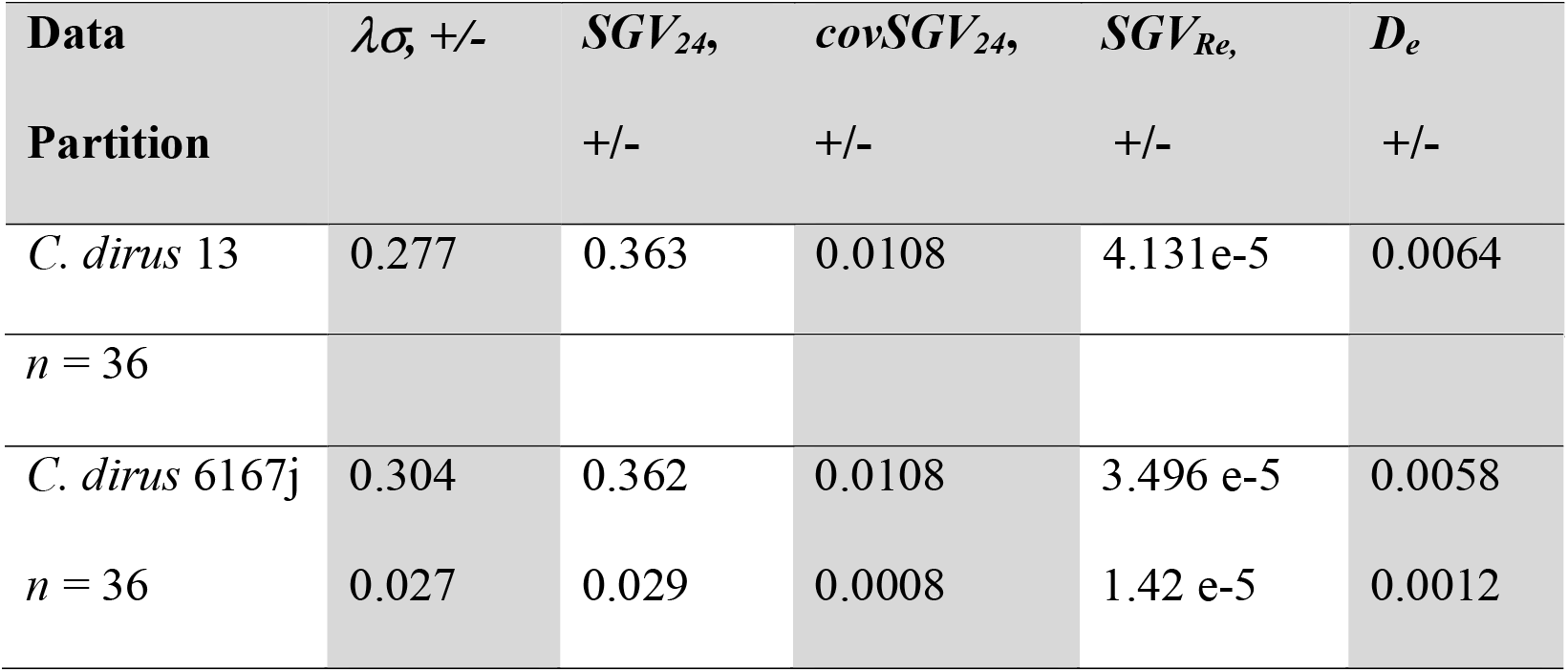
Whole jaw integration measures and other statistics for the *Canis dirus* mandible data sets. All reported confidence intervals are one standard deviation of 10000 jackknife or permutation replicates. The *D_e_* metric is significantly different between the two samples (*p* = 0.0049). Quantities listed are eigenvalue standard deviation (correlation matrix, 24 eigenvalues), (*λσ*); standardized generalized variance 24 (***SGV_24_***); standardized generalized variance 24 using the covariance matrix, and standardizing the eigenvalue vector to the sum of eigenvalues (***covSGV_24_***); standardized generalized variance effective rank (***SGV_Re_***); and effective dispersion (***D_e_***).

#### Effective rank-scaled SGV or ‘effective dispersion’

Calculation of the effective rank for the two populations indicates that the rank of Pit 61/67 is less than that of Pit 13 (Figure 4: *R_e_*, 13 > 61/67, *p* = 0.00148). To account for this difference in dimensionality dispersion, we modify the SGV to consider only the non-redundant eigenvalues of the covariance matrix. This metric includes only the eigenvalues that carry significant information, i.e. those up to the effective rank of the matrix. We label this quantity SGV_Re_, and calculate it as follows:

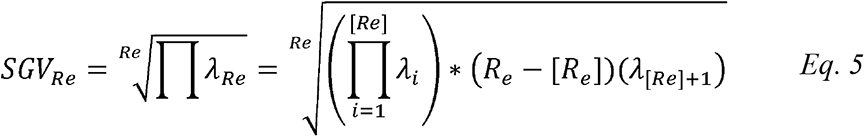

**Figure 4.**
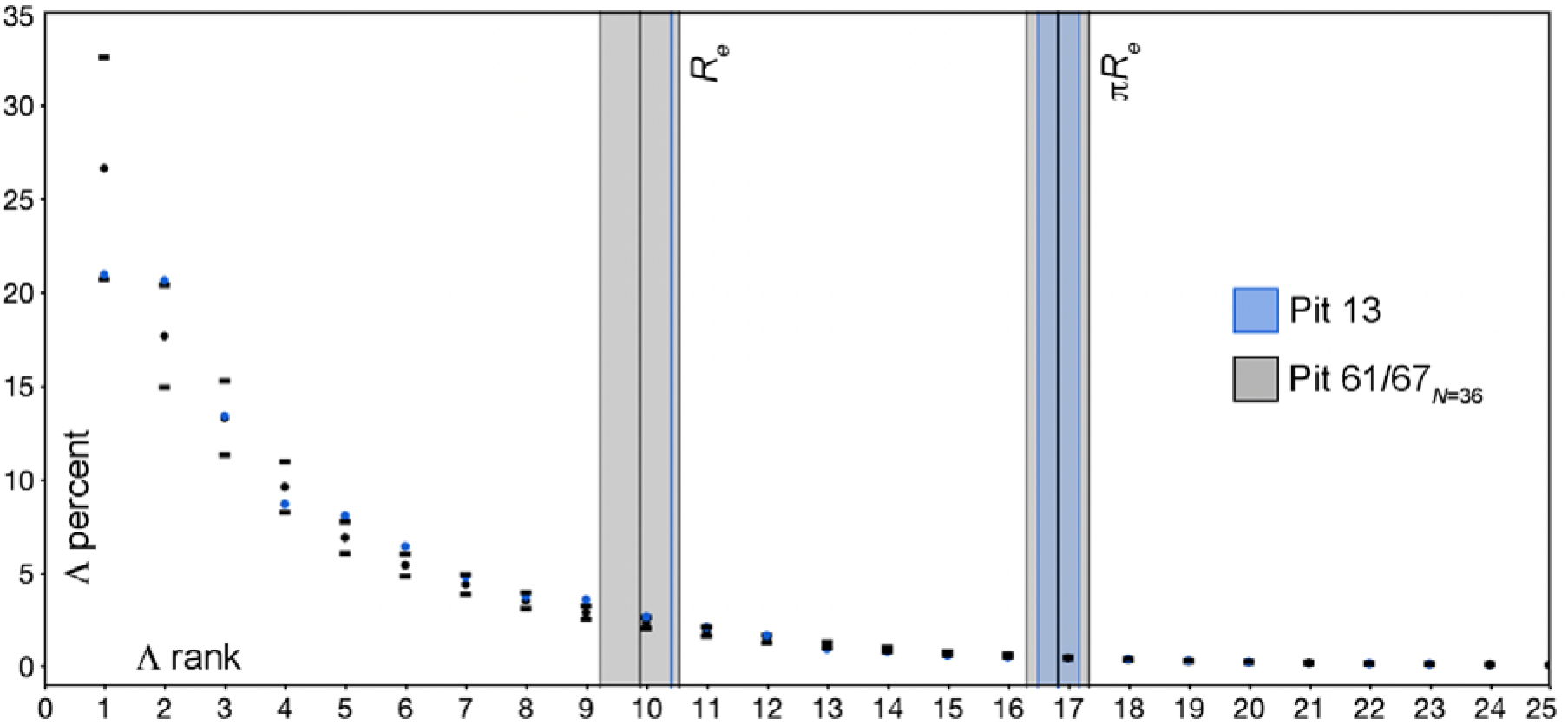
Scree plots and effective rank for GPA landmark data of *Canis dirus* jaws. Blue is Pit 13, black is Pit 61/67. Confidence intervals for effective rank (*R_e_*) are 10000 jackknife replicates of the Pit 61/67 sample to an *n* = 36. Eigenvalue dispersion appears greater in Pit 61/67 wolves, even though they have significant modularity and Pit 13 wolves do not. However the amount of total variance is greater in Pit 13, and the Pit 13 effective rank is significantly larger, so the variance is spread over a greater number of dimensions. The permuted dimensionality is also shown on both plots; this value is identical between pits (*πR_e_* = 16.84), demonstrating that the rank deficiency difference between the pits is due to phenotypic covariance only (i.e. the rank deficiency due to lack of matrix information is equivalent). The effective rank of both matrices is very deficient, with fewer than half of the 24 eigenvalues being meaningful. The effective rank of Pit 13 wolves is 10.414; the permuted effective rank is 16.841 +/− 0.345. The jackknifed effective rank for Pit 61/67 wolves is 9.89 +/− 0.656, while the permuted value at *n* = 36 is 16.832 +/− 0.519.

Where *R_e_* is the effective rank of the covariance matrix, and *[R_e_]* is the integer value of the effective rank. The SGV_Re_ is the *R_e_*^th^ root of the product of the eigenvalues up to the integer value of *R_e_*, multiplied by the decimal remainder of *R_e_* times the next smallest eigenvalue. The value for this statistic was calculated for the different data partitions using code written in R, with jackknifed confidence intervals for Pit 61/67 (Table 2; Appendix 1). Using this measure, Pit 13 wolves have significantly higher dispersion than those of Pit 61/67. Hence they are more integrated, while 61/67 wolves are more modular. The SGV_Re_ metric successfully recovers the increase in modularity exhibited by 61/67 wolves.

Previous authors rejected SGV as a statistic to measure dispersion because its distribution is not linear (Pavlicev et al., 2009); they prefered the standard deviation of the eigenvalues because of its linearity and because it has the same units as the input data. Because the SGV_Re_ is a measure of variance, one may utilize the definition of the standard deviation to transform the SGV_Re_ into a similar measure:

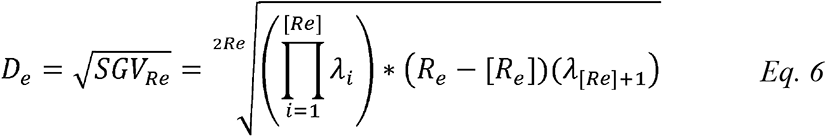

where the square root of the *SGV_Re_* is a new metric, *D_e_*, that we call ‘effective dispersion’. It measures dispersion in both variance and dimensionality together, and accounts for the rank deficiency that misleads the classical integration metrics. The *D_e_* metric is significantly different between the two samples (*p* = 0.0049; Table 2).

### Part 3: Matrix Information and Relative Dispersion

#### Permuted Rank Standardization

Sample size impacts effective rank; the effective rank of 61/67 wolves at *n* = 36 is 9.89, while the full matrix of *n* = 83 has an effective rank of 11.362 (Figures 2 and 4). Clearly the derivation of a version of *D_e_* that accounts for matrix size is desirable, as this will allow comparisons between matrices of different sizes. Pavlicev et al. (2009) accomplish a similar standardization for eigenvalue standard deviation by dividing the observed eigenvalue standard deviation by its maximum possible value to yield their “relative standard deviation”. Because they use the correlation coefficient in their calculations, the minimum possible correlation in a matrix is simply the number of traits minus one, because each trait adds an additional unit of uncorrelated variance. A relative version of effective dispersion is more difficult to calculate because the minimal covariance in a matrix is not derivable from first principles, because the input variance for each variable is not one. Also, because the matrix is rank deficient this must also be considered, suggesting that an approach based on effective rank is necessary. The quantity needed for standardization must therefore preserve the variances of the input variables, but remove their statistical covariance, and must also be treated for rank deficiency. One can generate a quantity with these properties by permuting the columns in **X** and then calculating the resulting covariance matrix. This matrix will preserve the input variances on the diagonal, but will remove statistical covariance on the off-diagonal (although covariances will still be non-zero in a rank-deficient matrix). The effective rank of the permuted matrix can then be calculated, yielding *πR_e_.* This permuted effective rank is the maximum matrix rank given the variable input variances and no statistical correlation among them. It is the amount of matrix information, and is driven by sample size in reasonably complex shapes. It can be used as a basis for standardizing *D_e_* for matrix size.

The appropriate scaling is based on consideration of what the effective dispersion actually is, namely the square root of the geometric mean of a subset of the eigenvalues of **K**, or alternatively, the root of the variance of one dimension of the hyperellipse. Because *D_e_* it is the square root of a mean eigenvalue (a variance), it has the same units as the original data (Pavlicev et al, 2009), and we would like to conserve this property in the scaled coefficient. Therefore there should be no variance term in the standardization on dimensional grounds. More importantly, the other property of *D_e_* is that it has been limited to its non-redundant information content, and we wish to scale *D_e_* to an expectation of maximum possible information content per axis. We are not concerned with scaling for the variance in the matrix, but for the amount of information in one dimension of an uncorrelated matrix that conserves this variance. The appropriate scaling term is therefore one rank of the permuted matrix, or 1/*πR_e_*. This yields the definition for the “relative dispersion”, or *D_r_*:

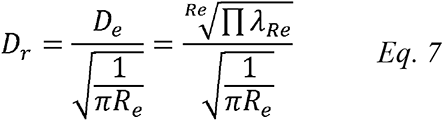

#### Relative Dispersion of *Canis dirus* and *Smilodon*

To demonstrate the use of relative dispersion, we turn to the third data set used in this paper, that of RLB *Smilodon fatalis* (Meachen et al., 2014). These data comprise 14 two-dimensional landmarks on 81 jaws, hence are quite similar to the dire wolf data (11 of the 14 landmarks are homologous between *Smilodon* and *Canis dirus*). We began by calculating the effective rank for *Smilodon* and comparing it to Pit 61/67 dire wolves (Figure 6). Both samples were bootstrapped for confidence intervals, and the sample size for 61/67 was held at 81 during the bootstrap. Both procedures lower effective sample size, and so reduce matrix information and hence rank. The exact values for integration statistics are reported in Table 4, along with the bootstrap means and standard deviations. Figure 6 plots the *n* = 81 bootstrap values for effective rank.

**Figure 6.**
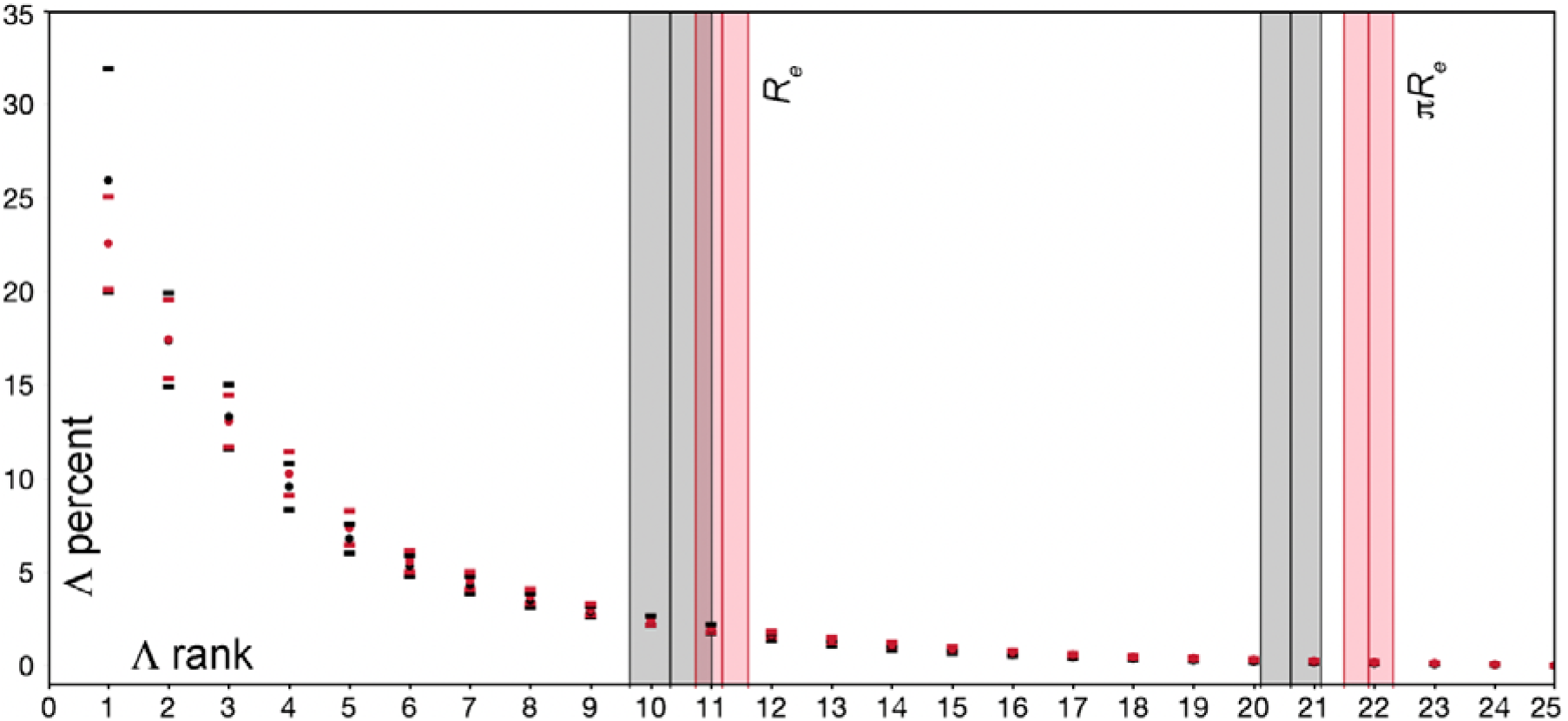
Scree plot comparison for *Canis dirus* and *Smilodon* data sets, 14 landmarks on jaws, at *n* = 81. Plotting conventions are the same as those used in Figure 2. *Canis dirus* 61/67 is shown in black, *Smilodon* in red. Both data sets have been bootstrapped for a confidence interval (10000 replicates), hence the values shown here are less than the precise values of *R_e_* for *C. dirus* (11.36) and *Smilodon* (12.32; Table 4). Because bootstrapping lowers the effective sample size, this drop in rank illustrates the strong sensitivity of the effective rank calculation to sample size. The plotted means and standard deviations are: *Canis dirus* 61/67, *R_e_* = 10.33 +/− 0.677; *πR_e_* = 20.614 +/−0.503. *Smilodon R_e_* = 11.184 +/−0.438; *πR_e_* = 21.91 +/− 0.503.

*Smilodon* and *Canis dirus* jaws are similar in that covariance matrices for both are highly overdetermined, although the effective rank of *Smilodon* is significantly greater. The permuted ranks of both taxa are much closer to 24 at a sample size of 81 compared to *Canis dirus* Pit 13, at a sample size of 36. This implies that the permuted rank of the matrices should converge toward 24 at large *n* (probably over 100, in accord with sample size requirements of other integration metrics as shown by Grabowski and Porto, 2017).

The relative dispersion is comparable among data sets of different sample sizes, as demonstrated in Table 4 (the *D_e_* of the 61/67 *n* = 36 jackknife sample is 0.0058, while that of the full *n* = 83 data set is 0.0052; the *D_r_* of both partitions is 0.0238). The values in Table 4 demonstrate that relative dispersion functions as intended; it is robust to difference is sample size, and successfully captures the modularity evolution between dire wolf populations. The comparison for *D_r_* is marginally significant, while that for sample-size controlled *D_e_* was strongly significant (Table 2); the reasons for this loss of power were not investigated. Lastly, the *D_r_* metric shows that the *Smilodon* jaw is much more tightly integrated than that of *Canis dirus*. We note that the *SGV_24_* successfully recovers this difference, while the eigenvalue standard deviation is still incorrect.

## Discussion

### Rank Deficiency and Its Treatment

In this paper we demonstrate that a typical GPA phenotypic covariance matrix is highly rank deficient. This fact is obvious from the scree plot, but quantifying the rank deficiency requires use of the Shannon, or information, entropy. Information entropy is used in signal processing to determine the amount a signal can be compressed without loss of information. We operationalize the Shannon entropy to a geometric morphometric context and use it to calculate that the information, or effective, rank of the dire wolf covariance matrix is less than half of the full rank. Geometric morphometric data sets have several characteristics that make rank deficiency more severe than in a matrix of linear measures. The first is the GPA procedure, which utilizes four degrees of freedom and hence removes four ranks from the covariance matrices dealt with here. This magnitude of this rank loss is known (Figure 1) and can be accommodated. By contrast, The magnitude of rank deficiency due to lack of matrix information is not known. We use a permutation approach to estimate this magnitude, allowing us to calculate the magnitudes of rank deficiency due to phenotypic covariance vs. the other sources. Phenotypic covariance accounts for about 9.6 ranks of deficiency in the dire wolf data set considered in Part 1, while lack of matrix information accounts for about 3 ranks, and the GPA accounts for 4 ranks. This non-phenotypic rank deficiency is worrisome, because misleads whole shape integration metrics that rely on the entire eigenvalue distribution.

To demonstrate the impact of rank deficiency on current integration metrics, we show that current whole shape integration metrics fail to identify the modularity change between dire wolf populations. Granted this is a minor change, from a one parameter model to a two parameter model, and the differences in whole-shape metric magnitudes are not large. However eigenvalue standard deviation is statistically significant in the wrong direction, while the *SGV_24_* seems to lack sensitivity. The *SGV_24_* does successfully capture the modularity difference between *Smilodon* and *Canis dirus*, where the integration difference is large. Yet eigenvalue standard deviation continues to fail. We also show that use of the correlation matrix leads to rank inflation in coordinate data; this rank inflation is noise, and the correlation matrix should not be used on raw coordinate data.

The failure of current integration metrics arises from the inclusion of a long tail of non-zero eigenvalues in the eigenvalue distribution. This tail of small values will have low dispersion, and eigenvalue standard deviation will be pushed lower as rank deficiency increases. The geometric mean of the eigenvalue distribution (the *SGV_24_*) will also be pushed lower, again due to the inclusion of many small eigenvalues. These facts are true on inspection but should be demonstrated; one immediate line of future research is a simulation study that quantifies the sensitivity of current integration metrics to information rank deficiency. This paper is meant as a theoretical development and proof of concept, and we do not attempt a simulation study here, although the behavior of the effective and relative dispersion metrics should also be explored by simulation. The seeming loss of statistical power between *D_e_* and *D_r_* deserves specific attention.

The use of the effective rank allows calculation of a modified form of the geometric mean of the eigenvalues, which is simply the mean of the non-redundant eigenvalues. We demonstrate that this new metric, *D_e_*, successfully recovers the evolution of increased modularity in Pit 61/67 wolves. However, matrix rank is very sensitive to sample size (Table 3), and we held sample size constant in Part 2. Therefore we introduce the concept of relative dispersion, *D_r_*, to account for differences in matrix information content in Part 3. Relative dispersion accounts for dispersion in dimensionality, as well as dispersion in variance, and it accounts for differences in matrix size by standardizing against the permuted information content. The metric *D_r_* should therefore be comparable among data sets, as we demonstrate via comparison with *Smilodon*. This comparison utilizes data sets with an equal number of landmarks (14); the sensitivity of relative dispersion to different landmark number requires quantification in a manner similar to the analysis of classical integration metrics by Grabowski and Porto (2017).

**Table 3.** Whole jaw integration measures for the *Canis dirus* jaw samples, and for the *Smilodon* data. The relative dispersion difference between Pit 13 and Pit 61/67 wolves is marginally significant (Welch’s *T* = 1.863, *p* = 0.068); Those between *Smilodon* and the two wolf data sets are highly significant (13, *T* = −10.85, *p* < 0.0001; 6167, *T* = − 17.29, *p* < 0.0001). The *Smilodon* jaw is much more tightly integrated than that of *Canis dirus*. Note that the *SGV_24_* also correctly captures the integration difference between *Smilodon* and *Canis dirus*, while the eigenvalue standard deviation still fails. Quantities listed are eigenvalue standard deviation (correlation matrix, 24 eigenvalues), (*λσ*); standardized generalized variance 24 (***SGV_24_***); standardized generalized variance effective rank (***SGV_Re_***); effective dispersion (***D_e_***); and relative dispersion (***D_r_***).

| Data Partition | $\lambda\sigma_{24}$ +/- | $SGV_{24}$ | $SGV_{Re}$ | $D_e$ | $D_r$ |
| --- | --- | --- | --- | --- | --- |
| *C. dirus* 13 $n = 36$ | 0.277 | 0.363 | 4.131e-5 | 0.0064 | 0.0263 |
| BS estimate | 0.316 | -- | 5.029e-5 | 0.00690 | 0.0283 |
| BS sd | 0.015 | -- | 2.283e-5 | 0.00162 | 0.0066 |
| *C. dirus* 61/67 $n = 83$ | 0.278 | 0.500 | 2.71e-5 | 0.0052 | 0.0238 |
| BS estimate | 0.296 | 0.421 | 3.373e-5 | 0.00567 | 0.0260 |
| BS sd | 0.025 | 0.030 | 1.311e-5 | 0.00112 | 0.0051 |
| *Smilodon* $n = 81$ | 0.250 | 0.575 | 7.20e-5 | 0.0085 | 0.0391 |
| BS estimate | 0.270 | 0.480 | 9.098e-5 | 0.00940 | 0.0432 |
| BS sd | 0.010 | 0.024 | 3.055e-5 | 0.00161 | 0.0074 |

### Dense Semilandmarks and Latent Dispersion

The relative dispersion metric accounts for information rank deficiency, but it is still space-specific. By this we mean that it is calculated in a shape space defined by the landmarks, not the entire shape, and the relation between these two spaces is not clear. For maximum utility, it is desirable that *D_r_* should be comparable among spaces defined by different, arbitrary sets of landmarks. In the case of two spaces defined by the same number of landmarks, as in the *Canis dirus* and *Smilodon* data compared here, one might suppose that relative dispersion is comparable even if the landmarks are not homologous (11 of the 14 landmarks are in fact homologous). Yet this is not true. A landmark-defined space with covariance matrix **K_V_** and an effective rank based on the Shannon entropy only captures the variance of the landmarks, not of the entire shape upon which those landmarks occur. One way to represent the total covariance of the entire shape, **K_T_**, would be to sample the shape with an arbitrarily large number of landmarks, so that as the magnitude of **V** increases **K_V_** would approach **K_T_**. Thought of in this way, a typical set of geometric morphometric data would comprise

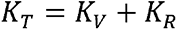

Where **K_R_** is the residual covariance in the shape not captured by the landmarks in **K_V_**. The residual matrix **K_R_** is of unknown magnitude in all mainstream applications of geometric morphometrics, and there is no guarantee that is negligible, nor that the proportion **K_R_/K_T_** is of constant magnitude among spaces, even if **K_V_**s are of equivalent effective rank. Therefore the relative dispersion metric defined above remains space-specific.

A recent approach to estimating residual shape variance is the employment of dense semilandmarks, where a curve or surface is first landmarked, and then densely sampled with an increasing number of semilandmarks until the coefficient of interest stabilizes (Marshall et al. 2019). This procedure can be employed here. Given a shape with variance **K_T_** and landmark variance **K_V_**, the residual covariance **K_R_** will decrease as **V** increases, so that

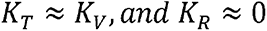

as the number of semilandmarks becomes large. The number of semilandmarks can be arbitrarily large, down to the pixel or voxel resolution of the image being digitized, but in practice semilandmarks need only be added until the coefficient stabilizes. Because the relative dispersion metric relies on information entropy, it will scale with the information added by each landmark, not by landmark number. Using this procedure allows characterization of the true, or latent, dispersion of the shape;

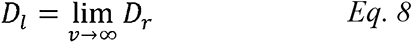

where the latent dispersion of the shape space, *D_l_*, is the relative disper sion of a matrix that is semilandmarked densely enough to asymptotically stabilize *D_r_*. Classical integration measures are not amenable to this procedure, because they consider all eigenvalues, and as the number of landmarks increases the rank of **K** will become increasingly rank deficient.

The latent dispersion metric should be useful for comparing shapes across arbitrary landmark spaces, and for assessing the fidelity with which the original landmark data capture phenotypic shape change. It is a general statement of the information content of a shape. Calculation of this metric would allow us to state definitively that the *Smilodon* data is tighter in shape dispersion than *Canis dirus*, assuming the observed pattern from 14 landmarks holds up. Yet given the rank deficiency in **K** demonstrated by the permuted matrices, we doubt there is sufficient residual variation not being captured by the landmarks to substantially change this result. *Smilodon* is a highly specialized hypercarnivore, and intuitively it should be more tightly adapted—and less complex— than a more generalized canid like *Canis dirus*, whose jaw should be more information rich.

## Conclusions

- Measures of whole-shape integration are reviewed; VanValen’s realization that dimensionality dispersion is a critical property is highlighted, and the Shannon, or information, entropy is employed to measure it as effective rank.
- Phenotypic covariance matrices for dire wolf jaws from two populations are introduced: Pit 13, deposited circa 19 kya, and Pit 61/67, deposited circa 14 kya. Both data matrices comprise X, Y coordinate data for 14 identical landmarks, after Procrustes superimposition.
- Use of the concept of information entropy allows identification of the magnitude and sources of rank deficiency in the Pit 61/67 sample. Rank deficiency is found to be high, with almost half not due to phenotypic covariance.
- Modularity model tests show that the only significant module is the ontogenetic one of the cheek teeth relative to the jaw corpus. This module is only significant in Pit 61/67 wolves; its absence in Pit 13 means that Pit 13 wolves are more integrated (best fit by a one parameter model), while 61/67 wolves are more modular (best fit by a two parameter model).
- Classical metrics of whole shape integration fail to recover the evolutionary increase in modularity in Pit 61/67 wolves. They fail because 1) they rely on the correlation matrix, which inflates trivial variance in geometric morphometric data; and 2) they measure only variance dispersion, while ignoring dimensionality dispersion.
- New metrics based on the Shannon entropy-modified SGV are defined to quantify the dispersion of phenotypic shapes spaces. The relative dispersion (*D_r_*) is a sample-size corrected version of the effective dispersion (*D_e_*); the latent dispersion is an asymptotic extension of *D_r_* to a dense semilandmark context. The relative dispersion is the core result of this paper; we restate it from Equation 7:

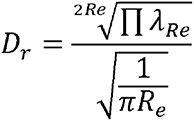
- New whole shape integration measures incorporating effective rank are successful at recovering the evolution of increased modularity in 61/67 wolves, and show promise for the characterization of phenotypic spaces among taxa.

## Supplementary Materials

Data Availability Statement: The data used in this paper are available as supplementary materials, and are also posted to the Dryad Digital Repository: http://dx.doi.org/10.5061/dryad.d7wm37q01

## Acknowledgements

We thank the curators and staff at the Tar Pit Museum, specifically John Harris, Emily Lindsey, Chris Shaw, Aisling Farrell, and Gary Takeuchi, for allowing us to collect data from the dire wolf specimens used in this project and for assistance of all kinds. We also thank Dr. H. David Sheets for his assistance using the IMP programs, including pcagen_7a. We thank the Graduate College for the Marshall University Summer Thesis Award and the Marshall University Drinko Research Fellowship to FRO for funding towards this project. Special thanks are due to Emily Lindsey, John Southon, Wendy Binder, and Larisa DeSantis for input on early versions of this manuscript, and to Bryan Carstens, Lauren Esposito, and two anonymous reviewers for thorough and very helpful input during the review process. Work by FRO was funded in part by NSF EAR-SGP 1757236 (Project SABER), and by a Drinko Distinguished Research Fellowship to FRO. We grudgingly thank Thom Yorke.

## Appendix 1

### R Code for Effective Dispersion Metrics, 1. Code for calculating permuted SVG, Effective Rank, and Effective Dispersion

~~~
#attach necessary libraries

library(dplyr)
library(broom)
library(readxl)

#Define a variable containing the GPA X,Y landmark data. The data is read in as a data frame from an Excel file.

GPALMData <- read_excel(“∼/Desktop/C dirus 14 LM/CDirus6167LM.xlsx”)

View(GPALMData)

EffRank=matrix(nrow=1,ncol=0)
SGV=matrix(nrow=1,ncol=0)
EffRankSGV=matrix(nrow=1,ncol=0)
EigenSumRec=matrix(nrow=1,ncol=0)

#Start permutation loop
for (i in 1:1000){

#Permute rows in each variable column vector independently for permutation test.
LMPrune = (GPALMData [3:30])
PermLM = apply(LMPrune, 2, sample, 83, replace=FALSE)
                                #sample size

#Define variable that is the covariance matrix of the above. Be sure to specify only the landmark columns in the data frame.
CVLM= cov(PermLM)
    #toggle for cov/cor

#Define object that is the eigenanalysis of the covariance matrix.
CVLMeigen=eigen(CVLM)

EigenVal<-CVLMeigen$values

#Define the summation of eigenvalues from the eigenanalysis.
EigenSum=sum(EigenVal)

#Create vector of scaled eigenvalues
PofK<-EigenVal/EigenSum

#Creat vector of lnPofK
lnPofK<-log(PofK)

Product<-lnPofK*PofK

SumProduct<-sum(Product, na.rm=TRUE)

ShannonEntropy <- -1*SumProduct

EffectiveRank<-exp(ShannonEntropy)

#Begin SGV24 calculation
EigenProduct<-prod(EigenVal[1:24])

StandGenVar<-(EigenProduct)^(1/24)

SGV<-cbind(SGV, StandGenVar)

#Begin ReSGV calculation
ERInteger<-as.integer(EffectiveRank)

EigenTrim<-EigenVal[1:ERInteger]

FractRankVar<-EigenVal[ERInteger+1]*(EffectiveRank-ERInteger)

ProductEffRankEigen<-prod(EigenTrim)*FractRankVar

EffectiveRankSGV<-(ProductEffRankEigen)^(1/EffectiveRank)

EffRank <-cbind(EffRank, EffectiveRank)

EffRankSGV <-cbind(EffRankSGV, EffectiveRankSGV)

EigenSumRec <-cbind(EigenSumRec, EigenSum)

cat(“\014”)

print(i)

}

#calculate and print st deviation and mean for each parameter

print(mean(EigenSumRec))
print (sd(EigenSumRec))

print(mean(EffRank))
print (sd(EffRank))

print (mean(SGV))
print (sd(SGV))

print (mean(EffRankSGV))
print (sd(EffRankSGV))
~~~

### R Code for Effective Dispersion Metrics, 2. Code for calculating SVG, Effective Rank, and Effective Dispersion

~~~
#attach necessary libraries

library(dplyr)
library(broom)
library(readxl)

#Define a variable containing the X,Y landmark data. The data is read in as a data frame from an Excel file.
GPALM <- read_excel(“∼/Desktop/C Dirus 14 LM/CDirus13LM.xlsx”) View(GPALM)

#initialize variables
EffRank=matrix(nrow=1,ncol=0)
SGV=matrix(nrow=1,ncol=0)
EffRankSGV=matrix(nrow=1,ncol=0)
EigenSumRec=matrix(nrow=1,ncol=0)
EffDispersion=matrix(nrow=1,ncol=0)

#Start bootstrap loop
for (i in 1:10000){
         bootsample=sample_n(GPALM, 36, replace = FALSE)
                                 #edit for n and T/F

#Define variable that is the covariance matrix of the above. Be sure to specify only the landmark columns in the data frame.
CVLM= cov(bootsample[3:30])
    #toggle cov/cor

#Define object that is the eigenanalysis of the covariance matrix.
CVLMeigen=eigen(CVLM)

EigenVal<-CVLMeigen$values

#Define the summation of eigenvalues from the eigenanalysis.
EigenSum=sum(EigenVal)

#Create vector of scaled eigenvalues
PofK<-EigenVal/EigenSum

#Creat vector of lnPofK and calculate effective rank
lnPofK<-log(PofK)

Product<-lnPofK*PofK

SumProduct<-sum(Product, na.rm=TRUE)

ShannonEntropy <- -1*SumProduct

EffectiveRank<-exp(ShannonEntropy)

#Begin SGV24 calculation
EigenProduct<-prod(EigenVal[1:24])

StandGenVar<-(EigenProduct)^(1/24)

SGV<-cbind(SGV, StandGenVar)

#Begin ReSGV calculation

ERInteger<-as.integer(EffectiveRank)

EigenTrim<-EigenVal[1:ERInteger]

FractRankVar<-EigenVal[ERInteger+1]*(EffectiveRank-ERInteger)

ProductEffRankEigen<-prod(EigenTrim)*FractRankVar

EffectiveRankSGV<-(ProductEffRankEigen)^(1/EffectiveRank)

#Make Effective Dispersion

EffDisperse<-sqrt(EffectiveRankSGV)

EffRank <-cbind(EffRank, EffectiveRank)

EffRankSGV <-cbind(EffRankSGV, EffectiveRankSGV)

EigenSumRec <-cbind(EigenSumRec, EigenSum)

EffDispersion <-cbind(EffDispersion, EffDisperse)

cat(“\014”)

print(i)

}

#calculate and print st deviation and mean for each parameter

print(mean(EigenSumRec))
print (sd(EigenSumRec))

print(mean(EffRank))
print (sd(EffRank))

print (mean(SGV))
print (sd(SGV))

print (mean(EffRankSGV))
print (sd(EffRankSGV))

print (mean(EffDispersion))
print (sd(EffDispersion))
~~~

### R Code for Effective Dispersion Metrics, 3. Code for calculating eigenvalue standard deviation

~~~
#attach required packages

library(dplyr)
library(broom)
library(readxl)

#Define a variable containing the X,Y landmark data. The data is read in as a data frame from an Excel file.

GPALM <- read_excel(“∼/Desktop/C dirus 14 LM/CDirus6167LM.xlsx”)
View(CDirus6167AllomResidLM)

BSSDEigen=matrix(nrow=1,ncol=0)

#Start bootstrap loop
for (i in 1:1000){
         bootsample=sample_n(GPALM, 83, replace = FALSE)
                                 #edit n, bootstrap or not

#Define variable that is the covariance matrix of the above. Be sure to specify only the landmark columns in the data frame.
CVLM= cor(bootsample[3:30])
    #toggle cov/cor

#Define object that is the eigenanalysis of the covariance matrix.
CVLMeigen=eigen(CVLM)

EigenVal<-CVLMeigen$values

EigenValTrim<-EigenVal[1:24]

EigenMean<-(sum(EigenValTrim))/24

EigenValTrimMean<-EigenValTrim/EigenMean

EigenK <- (abs(EigenValTrimMean-1))^2

Eigenvar <- (sum(EigenK))/24

SDEigen <- sqrt(Eigenvar)/4.7958

BSSDEigen <-cbind(BSSDEigen, SDEigen)

}

#calculate and pring st deviation and mean for each parameter

print(mean(BSSDEigen))
print (sd(BSSDEigen))
~~~

## Appendix 2

Testing the dire wolf data sets for modularity requires specifying candidate models. We use a heuristic approach based on the multivariate allometry vectors of interlandmark distances. Initial analysis consisted of calculating the interlandmark distances from the raw digitized landmark data **LM** (Figure A1). These distances were then evaluated for multivariate allometry by calculating the first principal component vector of the covariance matrix of *ln*-transformed distances; this procedure yields the multivariate allometry vector (Jolicoeur, 1963; O’Keefe et al., 1999; O’Keefe et al., 2013). Confidence intervals for the allometry vector coefficients where calculated by bootstrapping the input data (10000 replicates) in the R core package (R Core Team, 2014). The eigenvalues of the full 13-measure multivariate allometry vector were unusual, necessitating additional runs with some variables removed; all runs are reported in Table A1.

**Figure A1.**
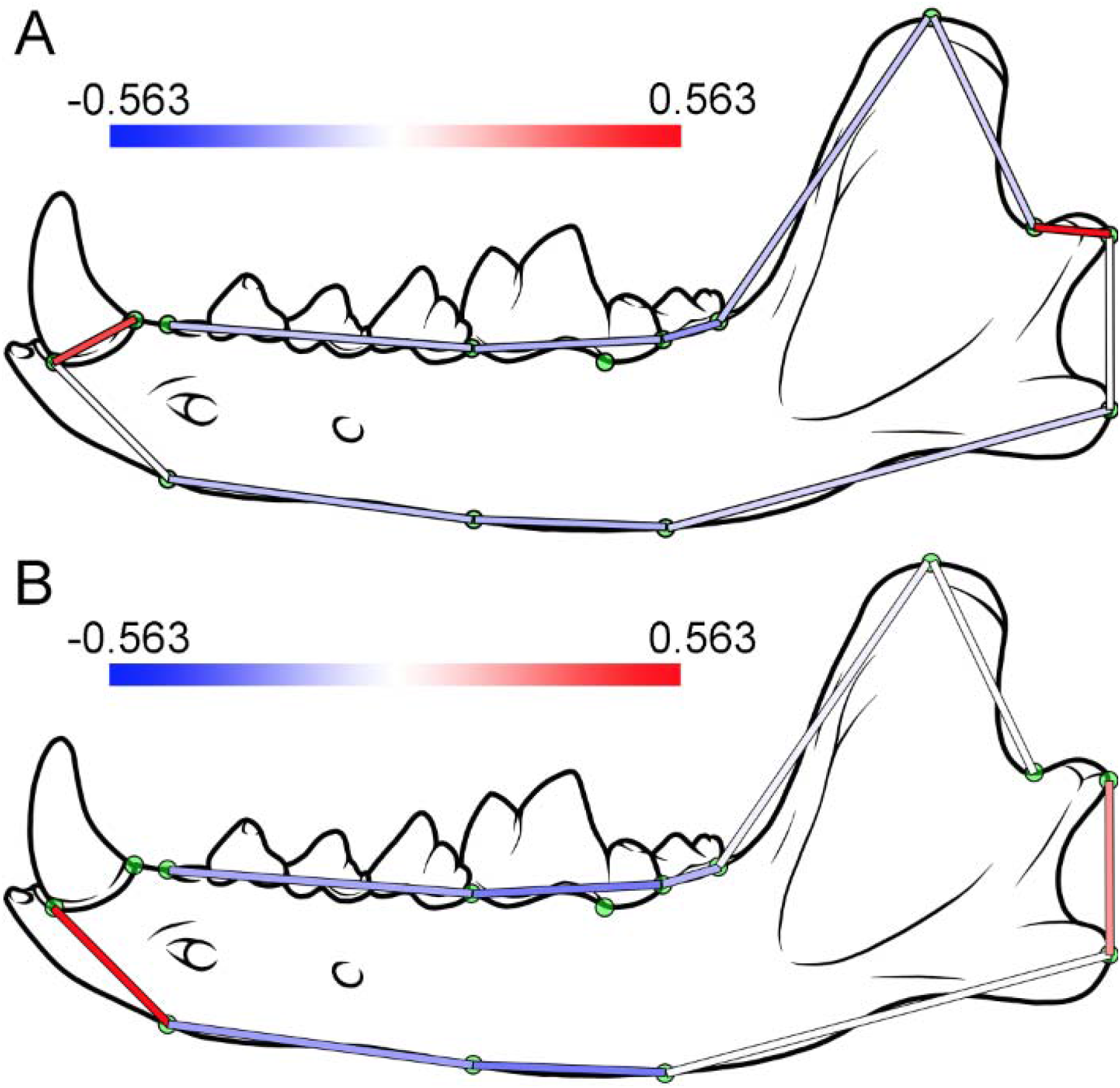
Interlandmark distances used in calculating multivariate allometry vectors. Redder lines indicate more positively allometric distances, blues lines more negative. Panel A depicts the 12-measure multivariate allometry vector, with each length heat mapped to its allometry coefficient from Table A1. Panel B shows the same for a 10- measure allometry vector. These different allometries suggest three different modules: the condoloid process and the canine, the check teeth, and the rest of the jaw.

**Table 1.** Allometry vectors calculated from interlandmark distances on *Canis dirus* dentaries as shown in Figure 1. Three vectors are shown; that for Pit 13, for Pit 61/67, and for the pooled sample. We also jackknifed the Pit 61/67 sample to establish a confidence interval; stars in this column indicate significant differences from Pit 13. The pooled vector was also bootstrapped, and stars in this column indicate differences from isometry in that vector. The isometric coefficient value for each vector is shown at bottom; confidence intervals for coefficient values are one standard deviation of 1000bootstrap replicates of each vector. Stars are 2 sigma. The heat maps in Figure 1A are indexed to the pooled coefficients.

| Distance | Pit 13 | Pit 61/67 | +/-, *n* = 36j | All | +/-, BS |
| --- | --- | --- | --- | --- | --- |
| 1-2 | 0.6543 | 0.3366 | 0.105** | 0.4889 | 0.056** |
| 3-4 | 0.0625 | 0.1503 | 0.036** | 0.1883 | 0.017** |
| 4-6 | 0.1339 | 0.1721 | 0.037 | 0.1500 | 0.022** |
| 6-7 | 0.1291 | 0.1561 | 0.071 | 0.1439 | 0.060** |
| 7-8 | 0.0706 | 0.2486 | 0.034** | 0.1927 | 0.020** |
| 8-9 | 0.0982 | 0.2457 | 0.038** | 0.2043 | 0.018** |
| 9-10 | 0.4183 | 0.4479 | 0.096 | 0.5682 | 0.043** |
| 10-11 | 0.2218 | 0.3494 | 0.056** | 0.2711 | 0.033 |
| 11-12 | 0.1140 | 0.2324 | 0.048** | 0.2105 | 0.023** |
| 12-13 | 0.1408 | 0.1562 | 0.041 | 0.1531 | 0.029** |
| 13-14 | 0.0593 | 0.1539 | 0.041** | 0.1842 | 0.012** |
| 14-1 | 0.5082 | 0.4540 | 0.133 | 0.2967 | 0.066 |
| Isometry | 0.289 | 0.289 |  | 0.289 |  |

The multivariate allometry vector for all 13 interlandmark distances is odd, with a single variable scoring very highly (the distance between the canine and the first premolar, landmarks 3-4), while the rest show significantly negative coefficients. This pattern indicates that the c1-p1 distance is highly variable, and that this variance is not correlated with any other variable. Further analysis shows that the c1-p1 distance crosses the boundary between two modules, accounting for its unique loading. We therefore discarded that distance and ran the analysis again with 12 distances. The resulting vectors for both populations are strongly positively allometric for two variables, the anteroposterior width of the canine (1-2) and the length of the condyloid process (9-10). Lengths associated with the other teeth are negatively allometric (Table A1). The covariation of canine width and condyloid process length suggests a functional unit, and provides one hypothesis for a testable module in the modularity tests. The lengths of the other teeth are all negatively allometric, as are the landmarks on the inferior margin on the jaw that are linked to tooth location. This finding of negative dental allometry in the cheek teeth has been documented before, both in *Canis lupus* (O’Keefe et al., 2016) and in *Canis dirus* (O’Keefe et al., 2014). Negative dental allometry is a consequence of the adult teeth forming early in ontogeny, erupting before the end of the first year, while full somatic growth is not attained until the end of the second year (O’Keefe et al., 2016). This negative dental allometry in the cheek teeth suggests a second module, comprising the lengths of the premolar arcade, the carnassial, and m2. We therefore specify three candidate modules for testing. They are the condyloid process and canine, the cheek teeth, and the rest of the jaw. The modularity models tested in Table 1 are permutations of these modules. We also note a significant allometric difference in the coronoid process in Pit 13 wolves; this is consistent with sexual diomorphism (O’Keefe et at., 2016), and we also tested modules including landmark 8 to test for this.

A valid criticism of our heuristic approach is that it does not test the model space exhaustively. The space of possible modules in a shape of 14 landmarks is 2^14^ – 1, or 16,383, small enough to allow exhaustive optimization using maximum likelihood. An alternative approach is weighted network analysis (Horvath, 2011), an effective method of model induction used with genomic data. We believe that the parts of the model space tested here capture all plausible candidate models, based on our analysis of the multivariate allometry vectors, and on subsidiary principal components analysis of the landmark data.

